# Poor concordance of floxed sequence recombination in single neural stem cells: Implications for cell autonomous studies

**DOI:** 10.1101/811455

**Authors:** Tyler Joseph Dause, Elizabeth Diana Kirby

## Abstract

To manipulate target gene function in specific adult cell populations, tamoxifen-dependent CreER^T2^ is widely used to drive inducible, site-specific recombination of LoxP flanked sequences. In studies of cell autonomous target gene function, it is common practice to combine these CreER^T2^-lox systems with a ubiquitously-expressed stop-floxed fluorescent reporter gene to identify single cells supposedly undergoing target gene recombination. Here, we studied the reliability of using Cre-induced recombination of one gene to predict recombination in another gene at the single cell level in adult hippocampal neural stem and progenitor cells. Using two separate stop-floxed reporters plus a Nestin promoter-driven CreER^T2^, we found that, in individual cells, expression of one reporter was a poor predictor of expression of the other. These findings imply that use of stop-floxed reporters to investigate cell autonomous gene function is likely to lead to false conclusions because recombination in separate genes shows poor concordance in individual cells.

## Introduction

In the adult mammalian brain, there are two primary neurogenic niches where neural stem and progenitor cells (NSPCs) proliferate throughout life: the subventricular zone (SVZ) and the dentate gyrus (DG) of the hippocampus (Gage, 2000). NSPCs in these regions create new neurons during adulthood, which incorporate into the preexisting circuitry in the olfactory bulb or DG, respectively, a process known as neurogenesis (Zhao et al., 2008). As evidence has mounted for the conservation of adult neurogenesis across species (Kempermann, 2016), understanding the functional properties of adult neurogenesis, as well as the molecular mechanisms that regulate it, has emerged as a major focus of research.

A common and powerful approach to dissecting molecular mediators of complex cellular processes like neurogenesis relies on gene ablation or over-expression to simulate gain or loss of function of target proteins. Genetic manipulation of adult neurogenesis requires cell-specific, inducible models that affect adult NSPCs selectively without influencing developmental counterparts. Ligand-dependent Cre recombinases, such as the tamoxifen(TAM)-dependent CreER^T2^, are widely-used for driving such inducible, site-specific recombination of LoxP flanked sequences in adult NSPCs (Feil et al., 2009; Semerci and Maletic-Savatic, 2016). Several CreER^T2^ mouse lines are currently available which use NSPC-specific promoters to drive Cre expression and therefore TAM-induced recombination in adult NSPCs (Semerci and Maletic-Savatic, 2016; Sun et al., 2014; Yang et al., 2015). In many studies, NSPC-targeted CreER^T2^-lox systems are combined with a ubiquitously-expressed stop-floxed fluorescent reporter gene to confirm recombination in NSPCs (Kirby et al., 2015; Tannenholz et al., 2016) (Sun et al., 2014). It is also common to use these fluorescent proteins more cell-specifically to investigate cell autonomous effects of recombination in the experimental gene of interest (Franz et al., 2019; Zhang et al. 2019; Zhou et al., 2018; Zimmermann et al., 2018). These cell autonomous experimental paradigms suppose that target gene and fluorescence reporter gene recombination occur in the same cell with high probability, allowing investigators to identify cell autonomous effects by comparing fluorescent and non-fluorescent cells.

To our knowledge, the assumption that fluorescent reporter expression reliably equates to recombination in a separate gene in the same cell has yet to be scrutinized. Within a cell, each recombination event is independent, and, with a transiently-activated Cre such as CreER^T2^, divergence of recombination status across loci seems probable. Here, we examine the reliability of using Cre-induced recombination of one gene to predict recombination in another gene at the single cell level in adult NSPCs. We found Cre-induced expression of one fluorescent reporter did not reliably predict expression of the other within a single cell *in vivo*. Given our results, we suggest that stop-floxed fluorescent reporters may misrepresent cell autonomous effects of gene recombination at a cell-specific level.

## Results

First, we explored the theoretical accuracy of using Cre-induced recombination of one gene to predict recombination in another gene. In this experimental paradigm, recombination of a stop-floxed fluorescent reporter (Gene R) is used as a marker of recombination in a target gene (Gene T) to make conclusions about the cell autonomous effect of target gene recombination (Fig 1A). This design assumes that presence of reporter protein indicates target gene recombination with high probability (a true positive signal) and absence of reporter protein indicates lack of target gene recombination with high probability (a true negative signal) (Fig 1B).

**Figure 1:**
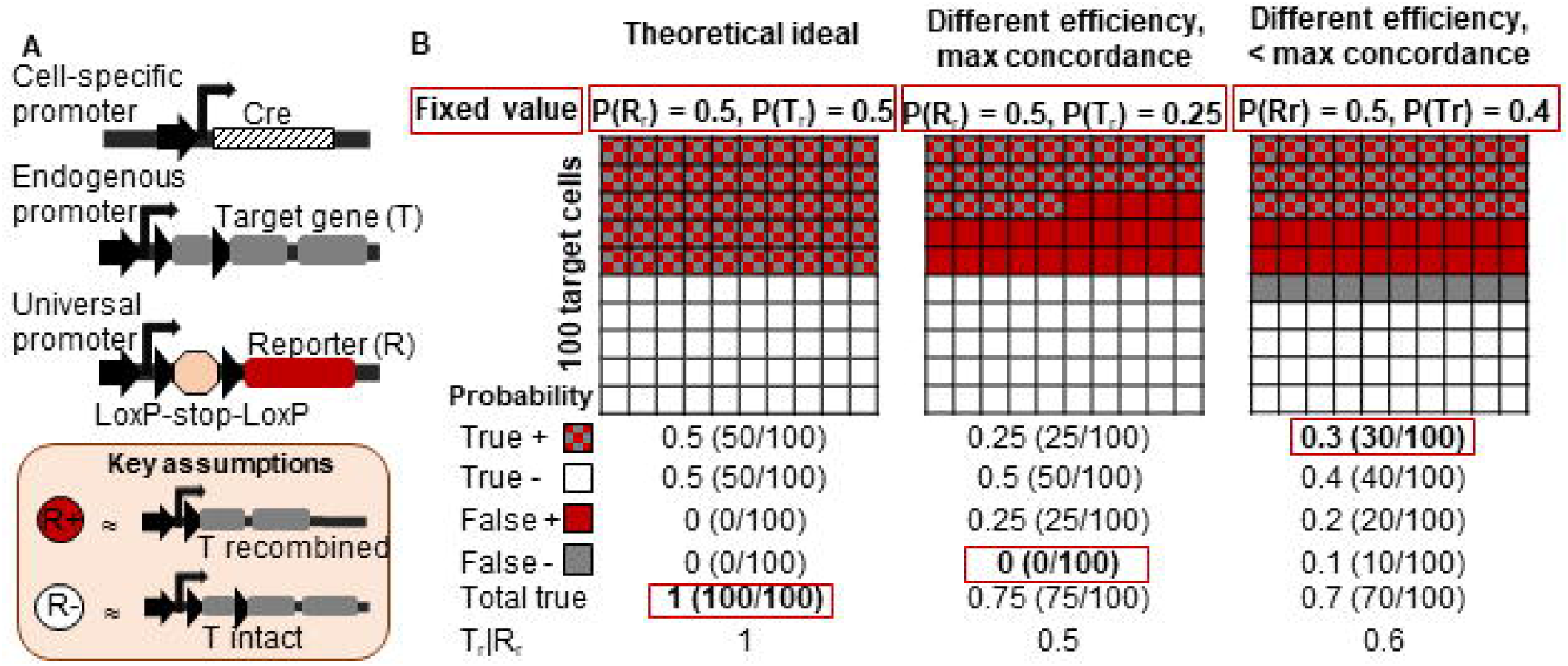
Theoretical probabilistic models reveal potential for error in a common paradigm for testing cell autonomous gene function. A) A generic schematic of the genetics used and assumptions employed in experiments where expression of reporter protein from a stop-floxed reporter construct is assumed to reliably indicate target gene recombination within a single cell is shown. B) The probabilities of true and false reporter signals are shown for 3 hypothetical scenarios with 100 target cells for visualization. The numbers in red boxes represent given values that are then used to determine the remaining probabilities. See also Supplementary Data sheet Figure S1.

To explore the consequences of deviations from the ideal of high true signal fraction, we performed probabilistic calculations of true signals based on several simplifying assumptions. First, we assumed recombination of gene T to be an all-or-none outcome (i.e. no possibility of heterozygosity). Next, we assumed that recombination does not occur in either gene when Cre is not expressed (i.e. minimal ectopic recombination). In many transgenic models, this assumption has been shown to hold true (Imayoshi et al., 2006; Kim et al., 2011; Yang et al., 2015). Third, we assumed that the probability of reporter recombination is larger than or approximately equal to the probability of target gene recombination ((P(R_r_) ≥ P(T_r_)). This assumption is based on the likely selection of stop-floxed fluorescent reporter constructs that recombine with high efficiency. Given these constraints, P(total true) = P(true+) + P(true-) = 1 + 2* P(R_r_) * P(T_r_|R_r_) – P(R_r_) – P(T_r_) (equation derivation in Sup Fig 1A).

In most experimental paradigms, P(R_r_) is known based on immunohistochemical quantification of percent of target cells expressing the reporter. For our estimates, we assumed an experimentally-feasible P(R_r_) = 0.5 (i.e. 50% of target cells showing recombination-dependent reporter expression). In the ideal outcome for this experiment, where total true signal is 100%, 50/100 cells would show simultaneous reporter expression and target gene recombination (Fig 1B, Sup Fig 1B, “Theoretical ideal”). However, while reporter gene recombination is frequently used as an estimate of target gene recombination, there is ample evidence that efficiency of Cre-LoxP recombination varies from gene to gene (Gray et al. 2017; Long and Rossi, 2009). We therefore examined the effect of setting P(R_r_) = 0.5 but P(T_r_) = 0.25. Using the best possible scenario where target gene recombination occurs only in reporter gene recombined cells (i.e. P(false -) = 0)), we found that P(T_r_|R_r_) = 0.5, meaning that identifying a cell as reporter positive only yields a 50% chance that the cell is also target gene recombined (Fig 1B, Sup Fig 1C, “Different efficiency, max concordance”).

We next estimated the effect of a less dramatic difference between P(R_r_) and P(T_r_) but when mixed with moderately suboptimal recombination concordance between the two genes within one cell. Using P(T_r_) = 0.4 and assuming that 75% of target recombined cells are also reporter recombined (i.e. P(T_r_)* 0.75 = P(true+)), we found that observing a reporter-expressing cell only yielded a 60% chance that that cell was also target gene recombined (i.e. P(T_r_|R_r_) = 0.6). Observing a reporter-negative cell in this case yielded a 10% chance that that cell was target gene recombined (P(false-) = 0.1) (Fig 1B, Sup Fig 1D, “Different efficiency, < max concordance”). These findings suggest that seemingly mild variation in recombination efficiency and suboptimal concordance rates for gene recombination could introduce substantial error in the method of using reporter expression as a marker of target gene recombination within a single cell.

To test the accuracy of using Cre-induced recombination of one gene to predict recombination of another gene in an in vivo experimental paradigm, we combined NestinCreER^T2^ mice with two commonly-used conditional reporter lines: Rosa-stop-floxed-EYFP (Srinivas et al., 2001) and Rosa-stop-floxed-tdTomato (Madisen et al., 2010) (Fig 2A). NestinCreER^T2^ mice express a tamoxifen (TAM)-sensitive Cre recombinase that drives recombination of floxed sequences in nestin-expressing NSPCs in the adult brain (Lagace et al., 2007). Both conditional reporters are inserted in the Rosa locus, a similarity intended to equalize both the Cre-accessibility of the two genes and the promoter activity driving their expression, thereby maximizing the likelihood of homozygous recombination within single Nestin-expressing cells after TAM exposure (Gray et al. 2017; Long and Rossi, 2009; Sandlesh et al., 2018).

**Figure 2.**
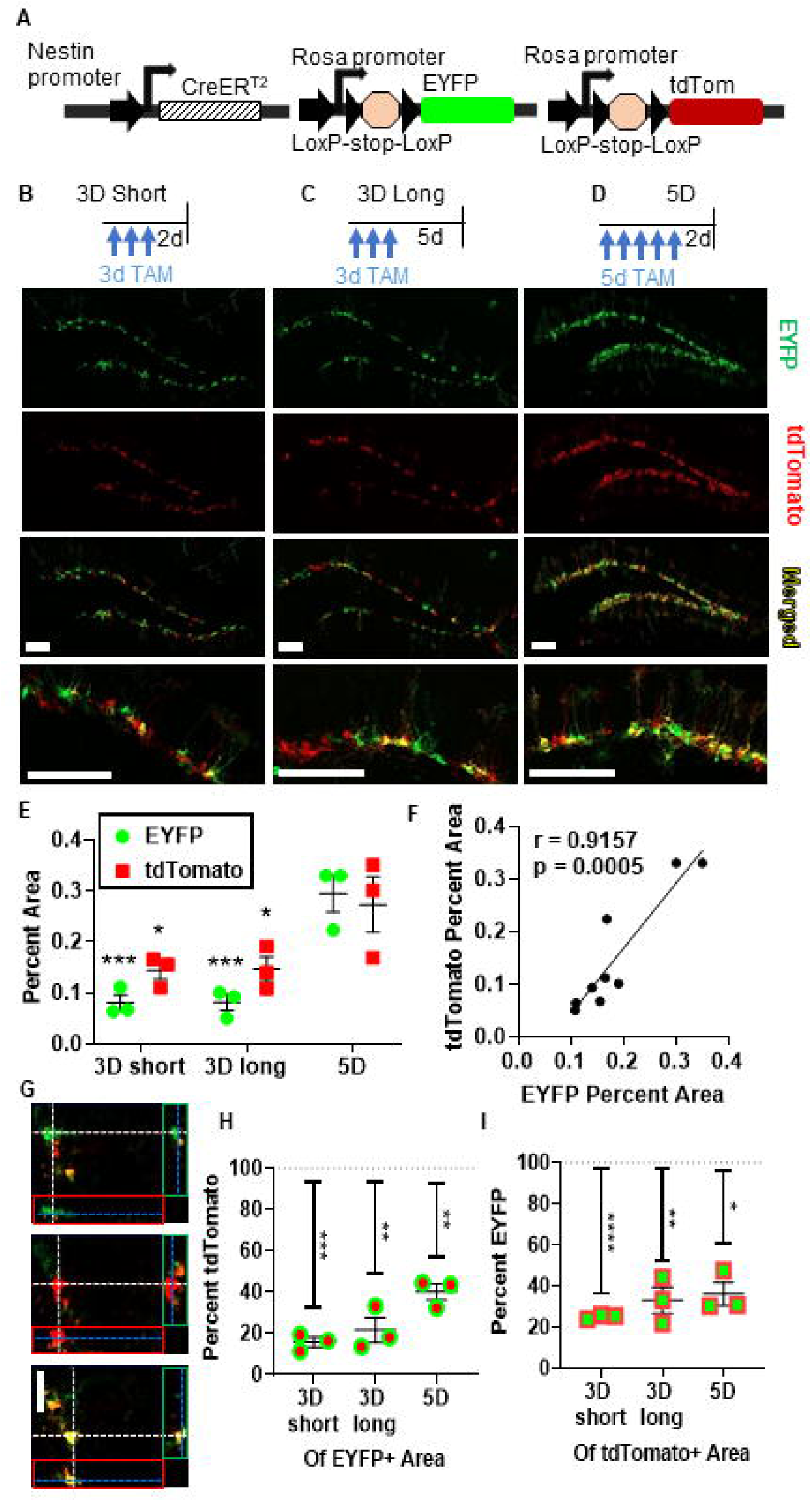
Recombination-dependent fluorescent reporter gene expression is correlated over the whole SGZ, but frequently fails to co-localize. A) A schematic of the transgenic mouse model employed in our experiments. B) Top, representative timeline of TAM injections and recovery for 3D short mice. Bottom, immunostaining of EYFP and tdTomato in SGZ. Scale bars 100 µm. C) Top, representative timeline of TAM injections and recovery for 3D long mice. Bottom, immunostaining of EYFP and tdTomato in SGZ. Scale bars 100 µm. D) Top, representative timeline of TAM injections and recovery for 5D mice. Bottom, immunostaining of EYFP, tdTomato in SGZ. Scale bars 100 µm. E) Comparison of EYFP+ and tdTomato+ DG percent area in 3D short, 3D long, and 5D mice. Tukey’s post hoc comparison within reporter type * p < 0.05, * * p < 0.01, * * * p < 0.001 vs 5D F) Correlation of EYFP+ and tdTomato+ DG percent area in all mice. R,p Pearson’s correlation. G) Orthogonal Images: Immunostaining and imaging of EYFP+, tdTomato+, and EYFP+tdTomato+ SGZ. Scale bar 20 µm. H) Comparison of EYFP+ colocalization in tdTomato+ area to theoretical 100% colocalization. I) Comparison of tdTomato+ colocalization in EYFP+ area to theoretical 100% colocalization. n = 3 mice per group. Data are shown as mean ± SEM. * p < 0.05, * * p < 0.01, * * * p < 0.001, * * * * p < 0.0001 one-sample t-test against a theoretical 100% (H,I).

First, to verify the TAM-dependency of reporter gene recombination in this model, adult NestinCreER^T2^;Rosa(EYFP/tdTom) mice were injected with oil (vehicle) or TAM once per day for 5 days. This TAM dosing schedule is widely-used and leads to efficient and specific recombination of floxed genetic sequences in NSPCs in adult NestinCreER^T2^ mice (Sun et al., 2014). As expected, oil-injected mice exhibited negligible ectopic recombination of either reporter gene (Sup Fig 1E) while TAM administration induced robust fluorescent reporter expression in DG NSPCs (Sup Fig 1F). These findings indicate that recombination of both reporter genes is tightly dependent upon TAM, as found in previous studies (Imayoshi et al., 2006; Kim et al., 2011; Yang et al., 2015).

We next examined the population-level efficiencies of recombination in both fluorescent reporter genes to determine if recombination frequency in one gene is a general predictor of recombination in a separate gene. To create a range of recombination rates, NestinCreER^T2^;Rosa(EYFP/tdTom) mice were submitted to 1 of 3 different TAM administration and recovery protocols: 3 days of TAM with 3 days of recovery (3D short), 3 days of TAM with 5 days of recovery (3D long) and 5 days of TAM with 3 days of recovery (5D) – the standard TAM regimen (Fig 2B-D). Analysis of the percent area occupied by fluorescent reporter proteins in the DG confirmed that a range of recombination rates was achieved, with 5D TAM leading to the highest level of reporter expression (Fig 2E). Across all three TAM groups, there was a strong and significant correlation between tdTomato and EYFP expression in the DG of TAM injected mice (Fig 2F). These findings suggest that mice with high recombination of one reporter gene also have high recombination of the other and support the possibility that recombination of one gene might be predictive of recombination in a second gene.

To examine reporter co-expression within single cells, we quantified EYFP and tdTomato fluorescence co-localization in 1 µm z-stack image series from adult DG. In these images, cells expressing EYFP alone appear green and cells expressing tdTomato alone appear red, while cells expressing both fluorescent reporters appear yellow (Fig 2G). Qualitative assessment of reporter expression revealed representation of each of these 3 possible colocalization phenotypes (Fig 2G). Using an automated co-localization tool to quantify reporter overlap, we found that EYFP+ percent area colocalization with tdTomato+ signal ranged from 15.84% in 3D short mice and 21.77% in 3D long mice to 36.71% in 5D mice, all of which were significantly less than 100% (Fig 2H). The converse—tdTomato+ percent area of EYFP+ area—showed slightly higher colocalization frequencies: 3D short 25.14%, 3D long 33.22%, and 5D 36.30%, though all still significantly less than 100% (Fig 2I). These results suggest that the fluorescent markers are not coexpressed in the same NSPCs with high frequency.

To quantify recombination frequencies within NSPC subpopulations in the subgranular zone (SGZ), we used immunofluorescent labeling for phenotypic markers to identify reporter expression in radial glia-like NSCs (RGLs) and intermediate progenitor cells (IPCs) (Fig 3A,B). RGLs were identified based on a cell body in the SGZ with a GFAP+ radial process extending through the granule cell layer (Lugert et al., 2010) (Fig 3A). IPCs were identified as Ki-67+ nuclei in the SGZ (Mandyam et al., 2007) (Fig 3B). Recombination frequency of each reporter gene individually was similar to that reported in previous studies using NestinCreER^T2^ mice (Sup Fig 2A-D). In addition, similar to our findings using fluorescent reporter area, recombination rates of EYFP and tdTomato reporter genes were positively associated (Sup Fig 2B,D). However, at the single cell level, we observed substantial populations of EYFP+ tdTomato- and tdTomato+ EYFP-RGLs and IPCs in all groups (Fig 3C,D,E,F), suggesting prevalent mis-match in recombination of the two genes in individual NSPCs.

**Figure 3.**
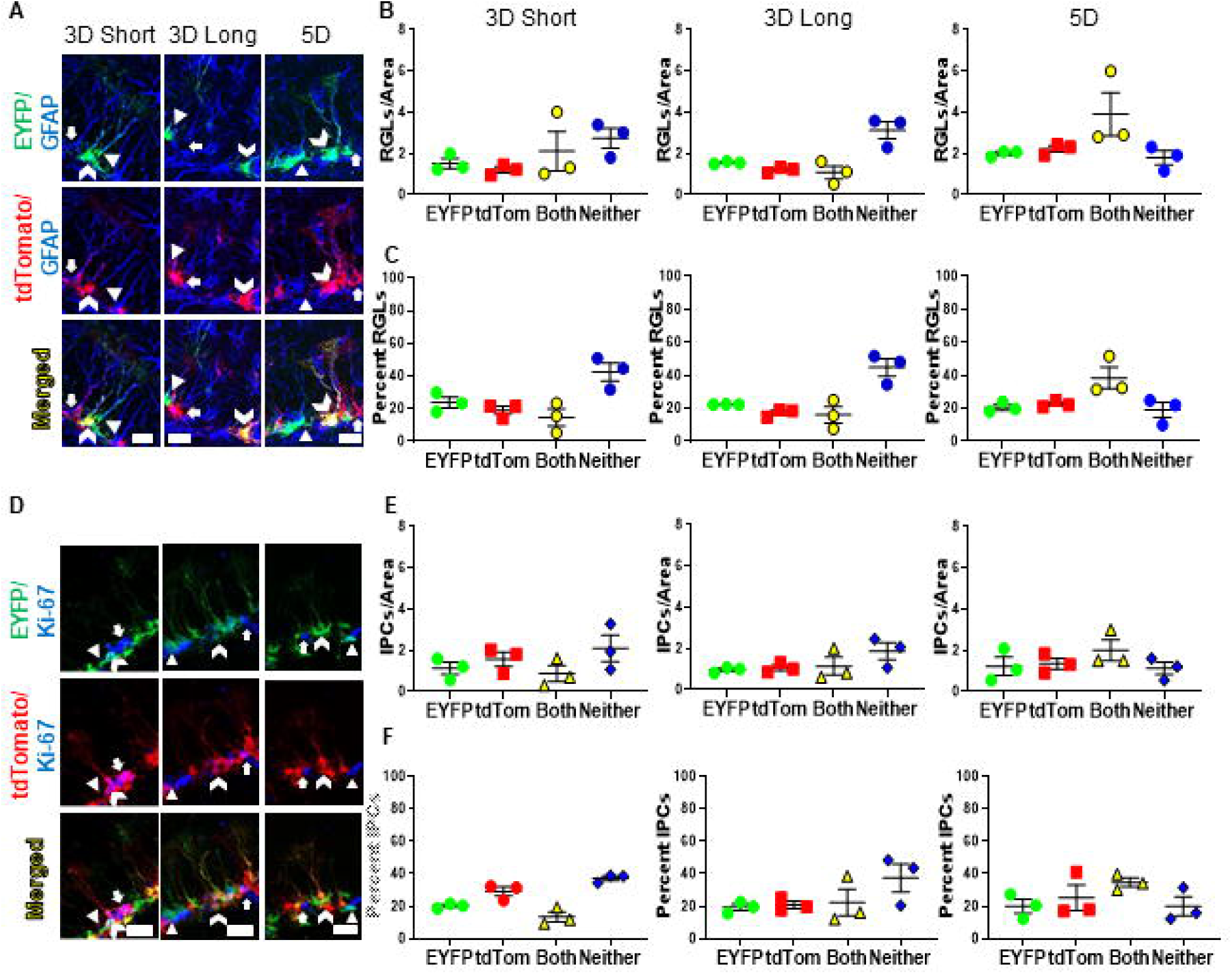
Recombination-dependent fluorescent reporter gene expression within RGLs and IPCs frequently fails to co-localize in the same cells. A) Immunostaining of EYFP, tdTomato and GFAP to identify SGZ RGLs. Scale bars 20 µm. Arrow head = EYFP+/tdTomato-RGL. Arrow = EYFP-/tdTomato+ RGL. Chevron = EYFP+/tdTomato+ RGL. B) Immunostaining of EYFP, tdTomato and Ki-67 to identify SGZ IPCs. Scale bars 20 µm. Arrow head = EYFP+/tdTomato-IPC. Arrow = EYFP-/tdTomato+ IPC. Chevron = EYFP+/tdTomato+ IPC. C) Density of SGZ GFAP+ RGLs co-expressing EYFP+/tdTomato-(EYFP), EYFP-/tdTomato+(tdTom), EYFP+/tdTomato+ (Both) and EYFP-/tdTomato-(Neither). D) Percentages of SGZ GFAP+ RGLs co-expressing EYFP, tdTom, Both or Neither. E) Density of SGZ Ki67+ IPCs co-expressing EYFP, tdTom, Both or Neither. F) Percentages of SGZ Ki67+ IPCs co-expressing EYFP, tdTom, Both or Neither n = 3 mice per group. Data are shown as mean ± SEM. Cells per area scale × 10^−4^. See also Supplementary Data sheet Figure S2.

To examine the consequences of the rates of single reporter recombination observed in our in vivo experiments, we applied the reporter expression frequencies observed in the 5D TAM group to a theoretical experiment where tdTomato (Td_r_, equivalent of R_r_ in Fig 2) is used to predict recombination in the EYFP reporter gene (E_r_, equivalent of T_r_ in Fig 2). In the 5D TAM group, the conditional probability P(E_r_|Td_r_) was 0.63 or 0.58 for RGLs and IPCs, respectively, meaning that there was a 63% or 58% probability that an RGL or IPC identified as tdTomato+ also had experienced recombination in the EYFP reporter gene (Fig 4A). The converse, P(E_r_’|Td_r_’), was 0.49 and 0.50 for RGLs and IPCs, respectively, meaning that identifying an RGL or IPC as tdTomato negative only yielded a 49% or 50% probability that the cell was also EYFP negative (Fig 4A). Comparing true positive and true negative signal in NSPCs across all 3 TAM protocols revealed that TAM protocol significantly interacted with the type of true signal (positive versus negative) (Sup Fig 3A, Fig 4B). Posthoc Tukey’s comparisons revealed that 5D TAM mice showed higher true positive signal than 3D short mice, but also lower true negative signal than 3D short or 3D long groups (Fig 4B). When true signal was summed, no difference was observed between TAM protocols (Fig 4C). Findings were similar when RGLs and IPCs were considered separately (Sup Fig 3B-E). These findings suggest that despite a wide range of recombination frequencies, none of the tested TAM protocols led better recombination concordance within single cells than the others.

**Fig 4.**
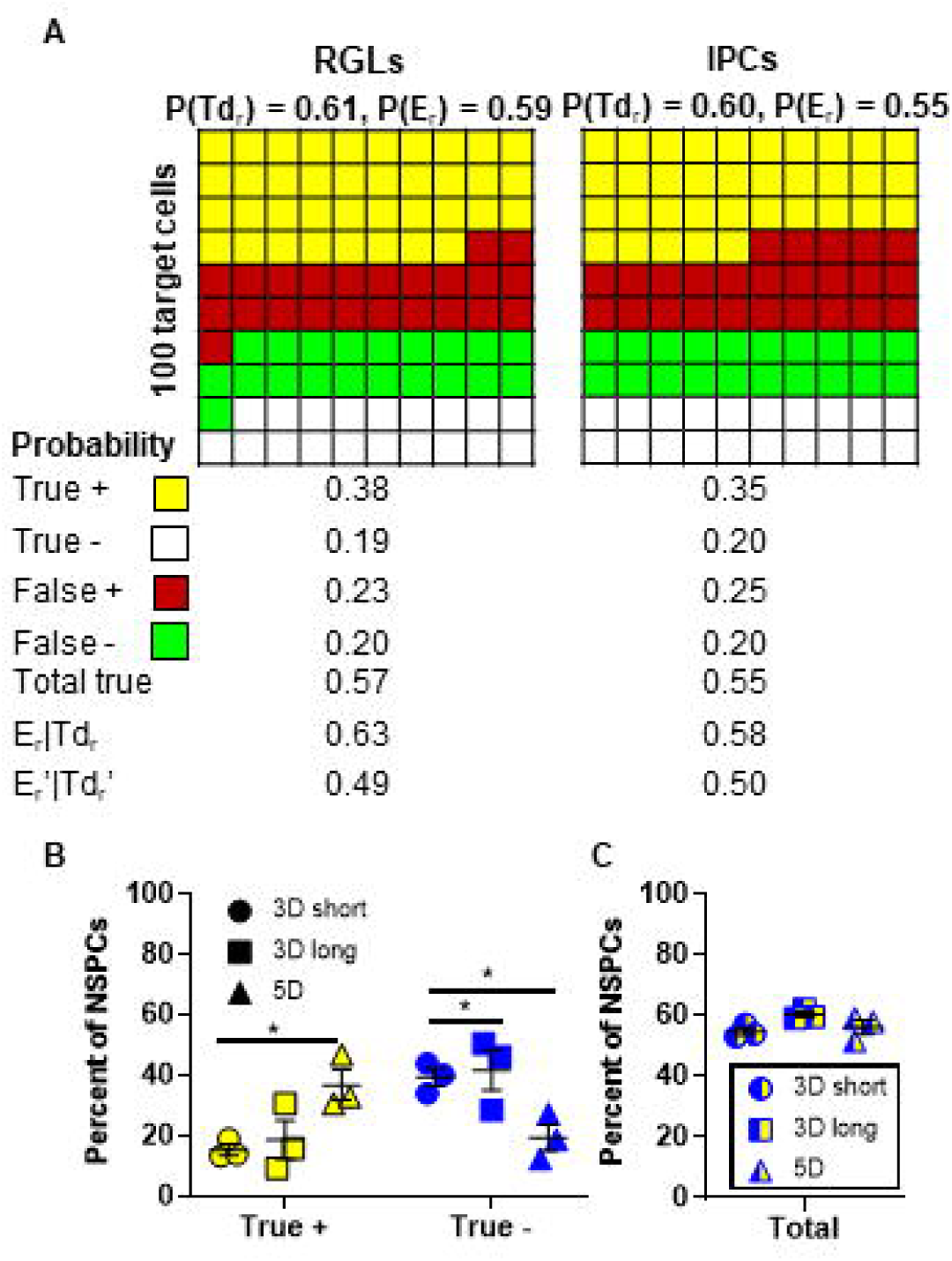
Using expression of one reporter to predict the other results in high false signals similarly across TAM protocols. A) The mean observed recombination frequencies from the 5D TAM group are represented here as reporter expression in 100 hypothetical target cells, either RGLs or IPCs. The probabilities of true and false signals are given as if tdTomato expression is being used to predict EYFP expression. B) The percent of RGLs and IPCs combined (total NSPCs) that show recombination in both reporter genes (true +) or neither (true -) is shown for 3 TAM groups. C) The total true signal between the 3 TAM groups is shown. n = 3 mice per group. Data shown are mean +/-SEM. * p < 0.05 determined by two-way ANOVA. See also Supplementary Data sheet Figure S3.

Though we focus on NSPCs in the SGZ, NSPCs are also found in the subventricular zone (SVZ). Similar to the SGZ, in the SVZ, TAM administration induced robust expression of both reporters that was correlated at the population level (Sup Fig 4A-B). However, as with the DG, we found significant divergence of expression at the single cell level. TdTomato+ colocalization within the EYFP+ area (57%) was significantly less than 100% in the SVZ, as was EYFP+ percent area of tdTomato+ area (40%) (Sup Fig 4C-D). These findings confirm and extend those from the SGZ by again showing that recombination of one fluorescent reporter gene does not accurately predict recombination of another at the single cell level in adult NSPCs.

## Discussion

Cell-specific gene manipulation is a powerful tool for investigating complex cellular mechanisms like neurogenesis *in vivo*. Here, we examine a common experimental model to assess cell autonomous function in which Cre-induced recombination of a stop-floxed fluorescent reporter is used to predict recombination of a target gene at the single cell level. The reliability of this model is contingent on the assumption that transiently-activated Cre induces independent recombination events equivalently within a single cell. We first examined this experimental paradigm with probabilistic calculations, which revealed that mild variation in recombination efficiencies and discordance between recombination in a fluorescent reporter gene and a target gene could lead to substantial error in identifying cells with target gene recombination. These data led us to examine the accuracy of this methodology in a mouse model. In our model, we quantified TAM-induced Cre-dependent recombination of two stop-floxed fluorescent reporter constructs in adult NSPCs. Our findings revealed that, while both reporters were highly expressed in NSPCs and global recombination rates of the two reporters was correlated, expression of either fluorescent reporter was a poor predictor of expression of the other in individual NSPCs. Based on our findings, we suggest that it is not reliable to assume that recombination of one gene is predictive of recombination of a second gene within single cells using TAM-sensitive Cre recombinases to drive recombination.

There is ample evidence that Cre-LoxP recombination varies from gene to gene, most likely due to differences in genomic accessibility to Cre enzyme (Gray et al. 2017; Long and Rossi, 2009). In our study, we used fluorescent reporters inserted in identical Rosa loci on separate chromosomes to minimize variation in Cre-accessibility. However, the vast majority of Cre-Lox models for testing cell-autonomous functions of a gene involve target genes in different loci than the stop-floxed reporter, adding an additional potential source of variance in recombination. We therefore hypothesize that our results represent a best-case scenario, and that real experimental paradigms likely have even poorer concordance between reporter expression and target recombination.

In adult neurogenesis studies, inducible gene manipulation is required to target cellular processes specifically in adulthood. However, activity of ligand-dependent Cre recominases within the nucleus are inherently limited and transient, reducing the likelihood of reaching 100% recombination efficiency across loci within a single cell. In non-inducible Cre-lox systems, in contrast, Cre is continuously present and active within cell nuclei, allowing for more exhaustive recombination of LoxP-flanked sequences. Lack of single cell concordance in LoxP recombination is therefore more likely to be a challenge in models using transiently inducible Cre than non-inducible Cre.

Our data suggest that to test cell autonomous gene function in adult NSPCs, it may be necessary to consider alternative models for identifying gene-manipulated cells. One paradigm that still uses inducible Cre would be to integrate fluorescent reporters in the target genomic sequence. Such models will likely require creation of new transgenic mice for many target genes, as few existing models include linked reporters in this fashion. An additional alternative method is to deliver transgenes plus cleavably-linked fluorescent reporters using viral vectors that target expression to specific cell populations based on viral serotype or cell-specific promoters. For example, certain adeno-associated viruses show preference for infecting specific NSPC subclasses and can be used to manipulate gene function in adulthood after stereotaxic delivery (Crowther et al., 2018).

In addition to tracking recombination in single cells, stop-floxed reporters are also frequently used to identify the cell population targeted by a Cre-driver line (i.e. population specificity), as well as overall recombination efficiencies. We found that both tdTomato and EYFP expression were similarly concentrated in adult NSPCs, suggesting that reporters can provide information about cell population specificity of recombination. We also found a strong correlation between recombination efficiency of the two reporters in individual mice, suggesting that reporters may reliably predict high-recombined subjects versus low-recombined subjects. However, it is unlikely that absolute recombination efficiencies can be extrapolated across genes due to the well-documented differences in LoxP recombination probability across genomic loci (Gray et al. 2017; Long and Rossi, 2009).

Our findings do not necessarily undermine previous studies, which have identified cell autonomous gene functions using stop-floxed reporters as a marker of target gene recombined cells (Franz et al., 2019; Zhang et al. 2019; Zhou et al., 2018; Zimmermann et al., 2018). If target gene recombination has a large cell-autonomous effect, it may still be detectable in reporter-expressing cells due to the subpopulation of cells in which fluorescent protein presence is a true signal. The frequent occurrence of false negative and false positive signals in this model is more likely to obscure smaller cell autonomous effects and lead to false negative findings when cell autonomous gene functions are in fact present.

In summary, our findings suggest that models of inducible gene manipulation combined with a ubiquitously-expressed stop-floxed fluorescent reporter are unreliable in their ability to identify cell autonomous effects at a single cell level *in vivo*. Future work is necessary to create reliable and cost-effective models that can be easily applied to the study of cell-autonomous effects across many target genes. Such models will be imperative for studying molecular mediators of complex cellular processes such as adult neurogenesis.

## Experimental Procedures

### Mice

All animal use was in accordance with institutional guidelines approved by the Ohio State University Institutional Animal Care and Use Committee.

### NSPC Identification and Manual Cell Counts

RGLs and IPCs were manually identified in 1 µm z-stack images using ImageJ then assessed for EYFP or tdTomato co-expression as described. Density of RGLs and IPCs was determined as the number of cells per area in the DG/SGZ.

### Statistical Analysis

Comparisons of more than two groups were performed using one or two-way ANOVAs followed by Tukey’s multiple comparisons. Correlations were performed using Pearson’s correlation. One-sample t-tests were used to compare difference from a theoretical value of 100%. All analyses were performed using Prism (v8.0; GraphPad Software) and p<0.05 was considered significant.

## Supporting information

Supplemental Statistics

## Acknowledgements

This work was supported by NSF grant IOS-1923094 to EDK. The authors report no conflicts of interest.

