## Supplemental Information for "Poor concordance of floxed sequence recombination in single neural stem cells: Implications for cell autonomous studies"

**Supplemental Figures, Titles and Legends**

**A**

P(total true) = P(true+) + P(true-)

Contraint: $P(R_r) \geq P(T_r$

P(true+) = P(Rr)*P(Tr|Rr)

P(true-) = 1- [P(true+) + P(false+) + P(false-)]

P(false+) = P(Rr) – P(true+)

P(false-) = P(Tr) – P(true+)

P(true-) = 1 – [P(true+) + P(Rr) – P(true+)
- P(Tr) – P(true+)]
= 1 + P(true+) - P(Rr) - P(Tr)
= 1 + P(Rr)*P(Tr|Rr) –P(Rr) – P(Tr)
= 1 + P(Rr)*(P(Tr|Rr)-1) – P(Tr)

P(total true) = P(Rr)*P(Tr|Rr) + 1 + P(Rr)*(P(Tr|Rr)-1) – P(Tr)

P(total true) = 1 + 2*P(Rr)*P(Tr|Rr) – P(Rr) – P(Tr)

**B**

**Theoretical ideal**

**Given:**

$P(R_r)$ = 0.5

$P(T_r)$ = 0.5

P(total true) = 1

P(total true) = 1 + 2*P(Rr)*P(Tr|Rr)
– P(Rr) – P(Tr)

1 = 1+ 2*(0.5)* P(Tr|Rr) – 0.5 – 0.5

**1 = P(Tr|Rr)**

P(true+) = P(Rr)*P(Tr|Rr)

P(true+) = 0.5*1

**P(true+) = 0.5**

P(true-) = P(total true) – P(true+)

P(true-) = 1 – 0.5

**P(true-) = 0.5**

P(false+) = P(Rr) – P(true+)

P(false+) = 0.5 -0.5

**P(false+) = 0**

P(false-) = P(Tr) – P(true+)

P(false-) = 0.5 -0.5

**P(false-) = 0**

**C**

**Different efficiency, max concurrence**

**Given:**

$P(R_r)$ = 0.5

$P(T_r)$ = 0.25

P(false-) = 0

P(true+) = P(Tr) - P(false-)

P(true+) = 0.25 – 0

**P(true+) = 0.25**

P(Tr|Rr) = P(true+) / P(Rr)

P(Tr|Rr) = 0.25/0.5

**P(Tr|Rr) = 0.5**

P(total true) = 1 + 2*P(Rr)*P(Tr|Rr)
– P(Rr) – P(Tr)

P(total true) = 1+ 2*0.5* 0.5 – 0.5 – 0.25

**P(total true) = 0.75**

P(true-) = P(total true) – P(true+)

P(true-) = 0.75 – 0.25

**P(true-) = 0.5**

P(false+) = P(Rr) – P(true+)

P(false+) = 0.5 -0.25

**P(false+) = 0.25**

**D**

**Different efficiency, < max concurrence**

**Given:**

$P(R_r)$ = 0.5

$P(T_r)$ = 0.4

P(true+) = 0.75 * $P(T_r)$ = 0.3

P(Tr|Rr) = P(true+) / P(Rr)

P(Tr|Rr) = 0.3 / 0.5

**P(Tr|Rr) = 0.6**

P(total true) = 1 + 2*P(Rr)*P(Tr|Rr)
– P(Rr) – P(Tr)

P(total true) = 1+ 2*0.5*0.6 – 0.5 – 0.4

**P(total true) = 0.7**

P(true-) = P(total true) – P(true+)

P(true-) = 0.7 – 0.3

**P(true-) = 0.4**

P(false+) = P(Rr) – P(true+)

P(false+) = 0.5 - 0.3

**P(false+) = 0.2**

P(false-) = P(Tr) – P(true+)

P(false-) = 0.4 - 0.3

**P(false-) = 0.1**

**E**

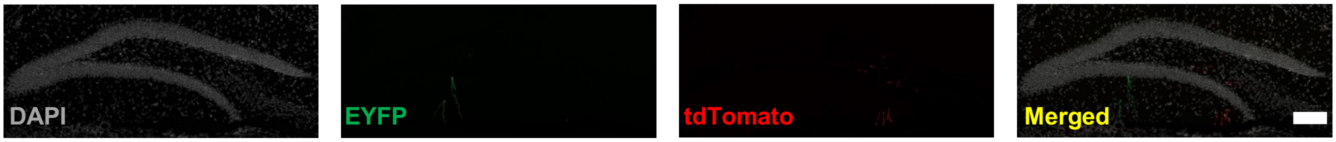


F

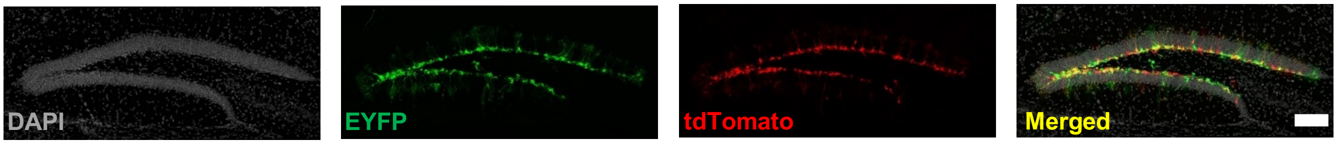

**Supplemental Fig 1: Equation derivation for true and false signal probabilities used for Fig 1. And comparison of Oil- and TAM-injected adult NestinCreER$^{T2}$;Rosa(EYFP/tdTom) mice.**

A) Assuming reporter recombination is greater than or equal to target gene recombination ($P(R_r) \geq P(T_r)$), probability of total true signal is derived using standard conditional and unconditional probability formulas.

B-D) Application of the equations in A to the scenarios described in Fig 1B are shown in full detail.

E) Immunostaining in the adult DG shows that oil administration does not stimulate expression of either reporter gene.

F) TAM administration induces robust recombination-dependent expression of both EYFP and tdTomato in the SGZ. Scale bars 100 μm.

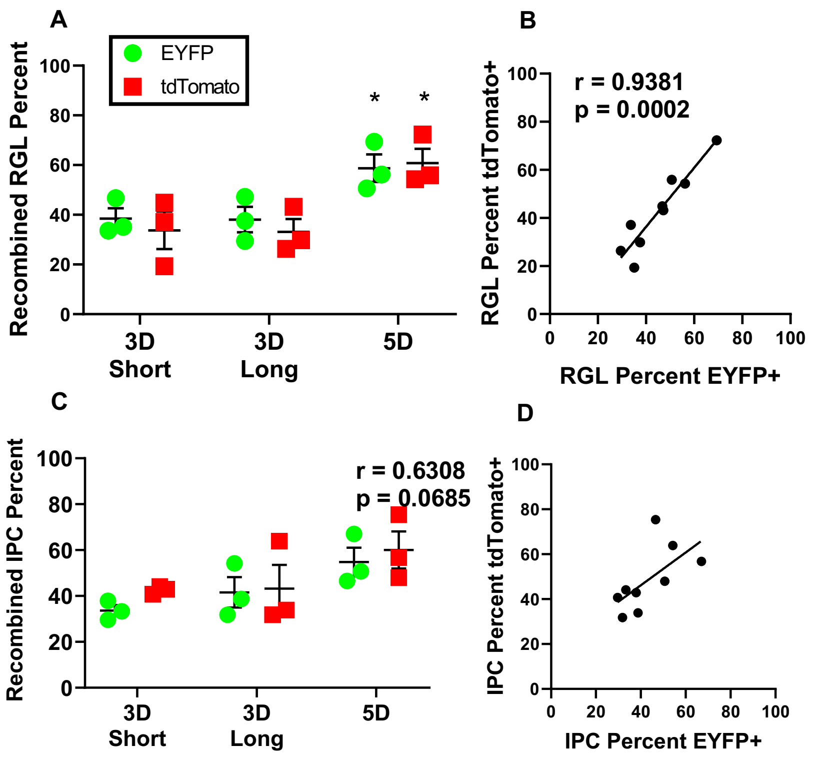


**Supplemental Figure 2. Supplemental information for Figure 3: Cell specific fluorescent reporter recombination frequency and correlation.**

A) Percent of EYFP+ or tdTomato+/GFAP+ RGLs in 3D short, 3D Long, and 5D mice.

B) Correlation of percent of GFAP+ RGLs that express EYFP and tdTomato in all mice.

C) Percent of EYFP+ or tdTomato+ IPCs in 3D short, 3D long, and 5D mice.

D) Correlation of percent of Ki67+ IPCs that express EYFP and tdTomato in all mice.

n = 3 mice per group. Data are shown as mean ± SEM. *$p < 0.05$, determined by two-way ANOVA (A,C) or Pearson's correlation (B,D).

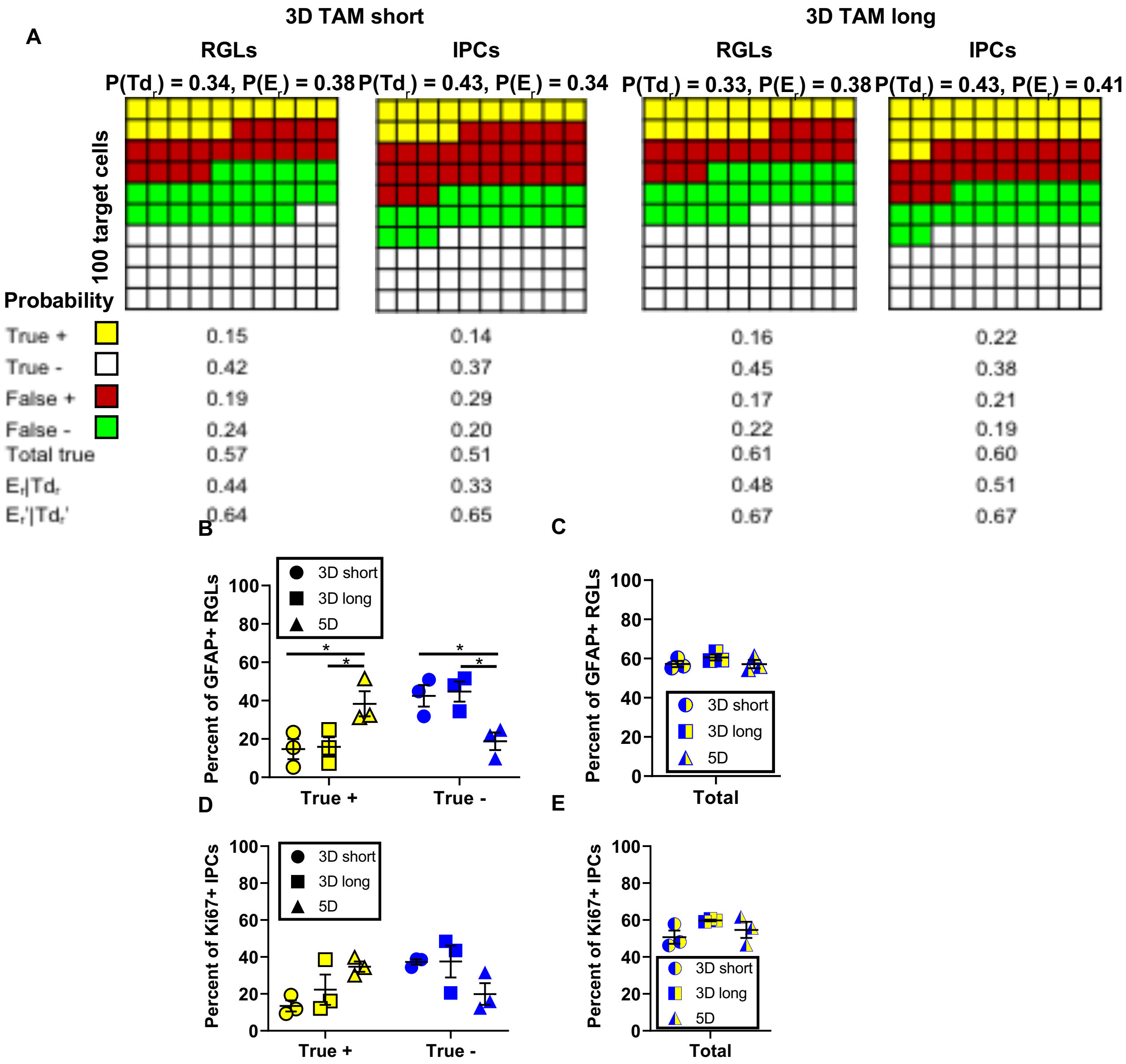


**Supplemental Fig 3. Supplemental information for Figure 4: Effects of cell type and Tam protocol of accuracy of using expression of one reporter to predict the other.**

A) The mean observed recombination frequencies from the 3D short (left) and 3D long (right) groups are represented here as reporter expression in 100 hypothetical target cells, either RGLs or IPCs. The probabilities of true and false signals are given as if tdTomato expression is being used to predict EYFP expression.

B) The percent of GFAP+ RGL cells that show recombination in both reporter genes (true +) or neither (true -) is shown for 3 TAM groups

C) The total true signal (+ and -) in GFAP+ RGL cells for the 3 TAM groups is shown.

D) The percent of Ki67+ IPCs that show recombination in both reporter genes (true +) or neither (true -) is shown for 3 TAM groups

E) The total true signal (+ and -) in Ki67+ IPCs for the 3 TAM groups is shown.

n = 3 mice per group. Data shown are mean +/- SEM. *p < 0.05, determined by two-way ANOVA.

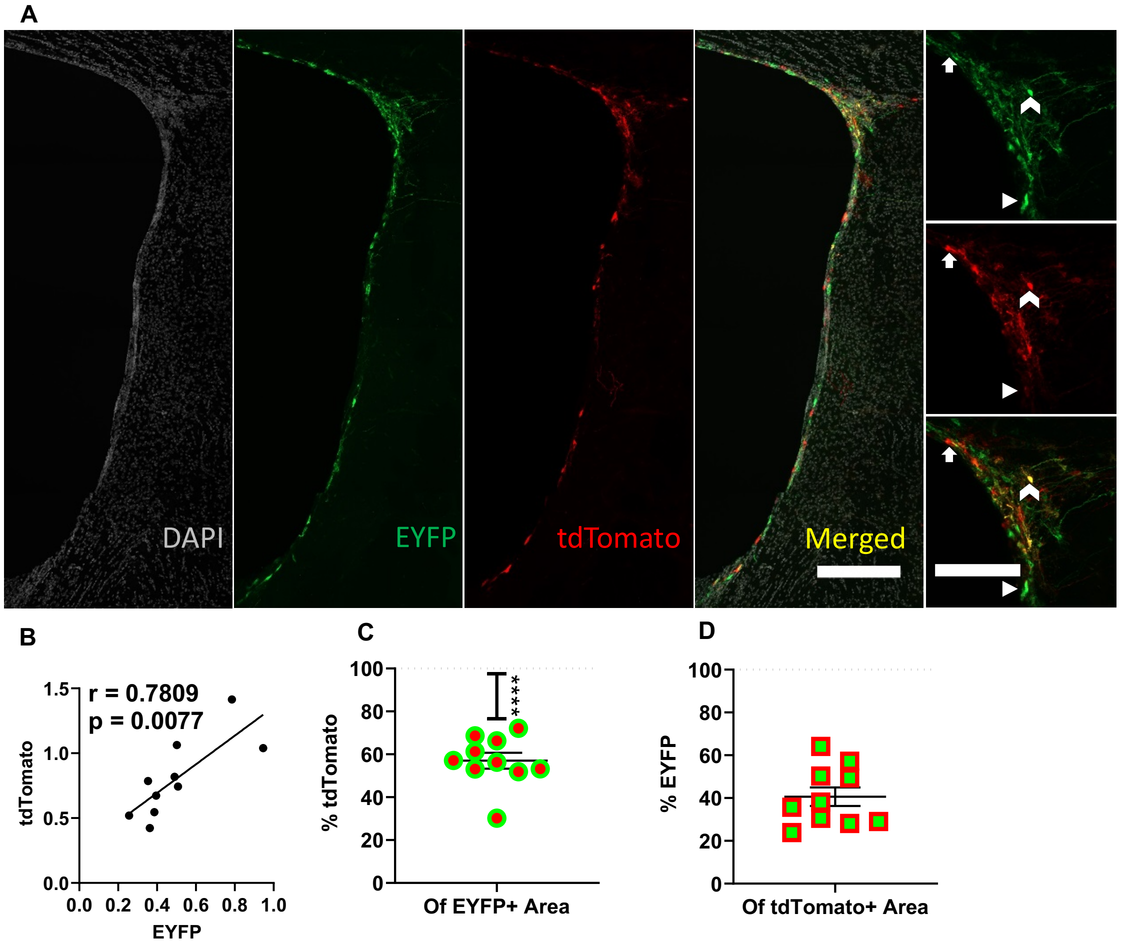


**Supplemental Figure 4. Fluorescent reporter recombination and colocalization in SVZ NSPCs.**

A) Immunostaining of EYFP, tdTomato in SVZ NSPCs. Scale bars 100 µm.

Arrowhead: EYFP+/tdTomato- NSPC

Arrow: EYFP-/tdTomato+ NSPC

Chevron: EYFP+/tdTomato+ NSPC

B) Correlation of EYFP+ and tdTomato+ SVZ percent area in all mice.

D) Comparison of EYFP+ colocalization in tdTomato+ area to theoretical 100% colocalization.

E) Comparison of tdTomato+ colocalization in EYFP+ area to theoretical 100% colocalization.

n = 9 mice. Data are shown as mean ± SEM. ****$p < 0.0001$ determined by Pearson's correlation (B) or one-sample t-test against a theoretical 100% (C,D).

### Supplemental Table 1: Primary and Secondary Antibody Table

| Primary Antibody | Vendor/ Product number | Dilution | Secondary | Vendor/ Product number | Dilution |
|---|---|---|---|---|---|
| Rabbit Anti-mCherry | Abcam ab167453 | 1:500 | Donkey anti-Rabbit IgG (H+L) Highly Cross-Adsorbed Secondary Antibody, Alexa Fluor 555 | ThermoFisher Scientific A-31572 | 1:500 |
| Hoescht | Fisher 33342 | 1:2000 | N/A | N/A | N/A |
| Goat Anti-GFP Antibody | Abcam Ab6673 | 1:1000 | Donkey anti-Goat IgG (H+L) Cross-Adsorbed Secondary Antibody, Alexa Fluor 488 | Fisher A-11055 | 1:500 |
| Mouse Anti-Glial Fibrillary Acidic Protein, Clone GA5 (GFAP) | EMD Millipore MAB360 | 1:1000 | Donkey anti-Mouse IgG (H+L) Highly Cross-Adsorbed Secondary Antibody, Alexa Fluor 647 | Fisher A-31571 | 1:500 |
| Rat Anti-Ki-57 Monoclonal Antibody | Invitrogen 14-5698-82 | 1:500 | Alexa Fluor® 647 AffiniPure Donkey Anti-Rat IgG (H+L) (712-605-153) | Jackson ImmmunoResearch 712-605-153 | 1:500 |

### Experimental Procedures

#### Mice

NestinCreER$^{T2}$ mice (Jackson #016261) were crossed with two conditional reporter lines: Rosa-stop-floxed-EYFP (Jackson #006148) and Rosa-stop-floxed-tdTomato (Jackson #007909). Mice were bred and maintained in the Ohio State University Psychology building mouse vivarium in standard ventilated cages on a 12h light cycle (lights on 0630h), with ad libitum access to food and water. All mice were 8-9 weeks old at the time of the experiment and housed in groups of 2-4.

#### Tamoxifen administration

TAM was dissolved in sterile sunflower oil at 20 mg/ml, overnight with agitation at 37 ˚C. TAM solution was stored at +4 ˚C for up to 1 week. TAM (or oil vehicle) was injected (180 mg/kg/d, IP) for 3d or 5d.

#### IF staining/antibodies

After 2 (3D short/ 5D mice) or 4d (3D long mice), mice were anesthetized with a 3% Ketamine Xylazine mixture then transcardially perfused ice-cold PBS. Harvested brain were fixed in 4% paraformaldehyde in 0.1 M phosphate buffer overnight at 4 deg C. After equilibration in 30% sucrose in 0.1 M phosphate buffered saline (PBS), 40 µm coronal brain sections were obtained in 1 in 12 series on a freezing microtome (Leica), and stored in cryoprotectant at -20°C until use. Brain sections were rinsed with PBS 3 times then incubated in a blocking solution containing 1% normal donkey serum (Jackson ImmunoResearch) and 0.3% Triton X-100 (Acros) in PBS. Sections were then incubated in primary antibody (Supplementary Table 1) diluted in blocking solution overnight at 4 deg C with rotation. The following day, after 3 rinses in PBS, cells were incubated in secondary antibodies (Supplementary Table 1) diluted 1:500 in blocking solution for 2 hours with rotation. The DG of the hippocampus was imaged in 15 µm Z-stacks at 20x magnification using a Zeiss Axio Observer Z.1 with apotome digital imaging system (Zeiss).

#### Automated Image Processing

Colocalization of immunofluorescent signal was analyzed with just another colocalization plugin (JACoP) software on ImageJ (Bolte and Cordelières, 2006). First, Z-stacks from each DG were separated into individual 1 µm thick images, 1 image per fluorescent channel. Images were thresholded then overlapped using anatomical features. The JACoP plugin then re-stacked the image files and analyzed overlap of EYFP and tdTomato. These data were used to determine EYFP and tdTomato percent area, the proportion of EYFP overlapping tdTomato and the proportion of tdTomato overlapping EYFP

#### NSPC Identification and Manual Cell Counts

RGLs were identified by their GFAP+ radial processes extending from the SGZ into the molecular layer, while cells with Ki-67+ cell bodies in the SGZ layer were identified as IPCs. The SGZ was defined as the zone spanning 2 cell body widths between the dense granular cell layer and the hilus. DG/SGZ area was drawn then measured in µm$^2$.
